# Gsmodutils: A python based framework for test-driven genome scale metabolic model development

**DOI:** 10.1101/430116

**Authors:** James P Gilbert, Nicole Pearcy, Rupert Norman, Thomas Millat, Klaus Winzer, John King, Charlie Hodgman, Nigel Minton, Jamie Twycross

**Affiliations:** Synthetic Biology Research Centre, University of Nottingham, Nottingham, NG7 2RD, United Kingdom; School of Biosciences, University of Nottingham, Sutton Bonington, Loughborough, LE12 5RD, United Kingdom; School of Mathematical Sciences, University of Nottingham, Nottingham, NG7 2RD, United Kingdom; Intelligent Modelling and Analysis group, School of Computer Science, University of Nottingham, Nottingham, NG8 1BB, United Kingdom

## Abstract

**Motivation:** Genome scale metabolic models (GSMMs) are increasingly important for systems biology and metabolic engineering research as they are capable of simulating complex steady-state behaviour. Constraints based models of this form can include thousands of reactions and metabolites, with many crucial pathways that only become activated in specific simulation settings. However, despite their widespread use, power and the availability of tools to aid with the construction and analysis of large scale models, little methodology is suggested for the continued management of curated large scale models. For example, when genome annotations are updated or new understanding regarding behaviour of is discovered, models often need to be altered to reflect this. This is quickly becoming an issue for industrial systems and synthetic biotechnology applications, which require good quality reusable models integral to the design, build and test cycle.

**Results:** As part of an ongoing effort to improve genome scale metabolic analysis, we have developed a test-driven development methodology for the continuous integration of validation data from different sources. Contributing to the open source technology based around COBRApy, we have developed the *gsmodutils* modelling framework placing an emphasis on test-driven design of models through defined test cases. Crucially, different conditions are configurable allowing users to examine how different designs or curation impact a wide range of system behaviours, minimising error between model versions.

**Availability:** The software framework described within this paper is open source and freely available from http://github.com/SBRCNottingham/gsmodutils

## 1 Introduction

Stoichiometric constraints based modelling for biological systems has been a mainstay of systems biology for several decades (*1, 2*). Given its flexibility, low barrier to entry and requirement only on minimal knowledge regarding the stoichiometry of metabolic networks this structural approach has become an extremely popular method for modelling steady-state behaviour of large, biochemical networks (*3*). Such large scale reconstructions are often referred to as genome scale metabolic models (GSMMs), as the processes is significantly aided through the advent of relatively inexpensive genome sequencing (*4, 5*). Indeed, owing to their ability to model complex aspects of metabolism, GSMMs have been widely adopted as a standard to elucidate and optimise of industrial biotechnology processes (*6*).

The reconstruction of GSMMs is an arduous process that follows a complex protocol to ensure model validity (*7*). Whilst many popular automated methods exists to construct GSMMs from reference genomes (*8, 9*), there is a still a significant amount of manual curation. However, treating the creation of models as an isolated “one-off” event ignores the significant amount of curation that is required for applications such as biotechnology.

As a consequence, a significant amount of work has gone into the management of genome scale models. The BiGG models database (*10*), for example, exists to provide a standardised repository of validated models that can be shared and reused. However, little focus is placed upon the collaborative design aspect of such models with few mechanisms existing to capture the differences between two model versions, *model deltas*. Similarly, the MetaNetX (*11*) system exists to provide a standardised namespace and toolchain for GSM analysis. However, such tools often make it difficult to understand the design decisions made by the initial model authors.

Furthermore, as with many areas of bioinformatic study the number available computational tools has become vast. This covers huge variety of software platforms including the COBRA toolbox for Matlab (*12*), ScrumPy and COBRApy in python (*9, 13*) with additional tools and libraries such as cameo (*14*), OptFlux in Java (*15*) and SurreyFBA (*16*). Whilst most of these tools are Open Source and follow standards, such as SBML (*17*), it is often challenging to replicate the initial modelling efforts conducted by authors of papers. Consequently, we feel that software tools are urgently needed to address this issue.

Similarly, The archetype design, build and test cycle of synthetic biology heavily relies on the usage of bioinformatics software and modelling to improve the production of natural products (*18*). In order to speed up the use of bioinformatics tools to produce high value platform chemicals, genome scale models are often used to discover methods for process optimisation *in silico*.

For example, many tools such as RetroPath (*19*), XTMS (*20*) and GEM-Path (*21*) suggest thousands of potential heterologous pathways. Many of these tools significantly increase the value of genome scale models, for example by coupling commodity production to an organism’s growth (*22*). These tools all suggest major changes to wild-type strains must be tracked and compared to allow models to remain relevant. In effect, mechanisms are required to relate to modified test and production strains.

Similarly, many conventional applications of genome scale models in systems biology have often suffered from unnecessary replication of work due to a lack of adherence to standards (*23*). For example, there are now many independently developed models of *Clostridium acetobutylicum* (*24, 25, 26, 27, 28*), an organism used in the production of solvents for around a century (*29, 30*). These models all exist to solve similar biological problems some being updates to the initial base models. However, there has been disagreement over fundamental biochemical properties of this anaerobic organism, notably with the focus on redox balancing (*27*). Such models also include updates based on improved genome annotations and the inclusion of fluxomic, transcriptomic and metabolomic characterisations (*28*).

Unfortunately, many of the results reported in (*24, 25, 26, 27, 28*) are difficult to compare or reproduce as the result of a number of issues. Often, the models lack a standard-isation of identifiers for reaction names, which makes direct comparison of model structure as well as differences between reactions a challenge. Where models are shared, it is often in non-standard spreadsheet formats, rather than SBML models. Indeed, even in the case of valid SBML models being made available at the time of publication few details are given as to how to run such models for conditions discussed in original articles.

In this paper, we present a software framework geared toward *test-driven* genome scale model development, a concept that is taken directly from good software development practices (*31*). By this we mean the notion that, as a model is curated to represent biological phenomena, much of the validation can be turned into specific test cases that can be repeated between model versions. We provide an example test case for *Clostridium autoethanogenum*, an organism that has had considerable focus in terms of genome scale models and how a working methodology using the software presented here can reduce repetition of work and improve the reproducibility of results. This article aims to summarise the main objectives of the *gsmodutils* software and we refer the reader to the software user guide for a more detailed exploration features.

## 2 Improving the design phase of industrial biotechnology

The design phase of industrial synthetic biotechnology, due to historical and regulatory convention, often follows a waterfall model for development of genome scale models. This is borne out of a number significant constraints in the development of genome scale models. The waterfall approach for modelling is characterised in Figure 1 where requirements for a model are captured at an early stage and are hard to refine during the production process.

In contrast, an *agile* methodology for the development of models places the focus on adapting work to new requirements (*31*). Such an approach best fits genome scale models because they are rarely created to investigate individual processes and, instead, capture the complexity of large systems. Genome scale models are intrinsically related to available genome annotations. Such annotations rely heavily on automated matches to related species, with the characterisations of individual genes or changes in cofactors and substrates for specific reactions often being left to a few of critical interest (*32*). This modelling formation costs an in depth understanding of dynamic behaviour. However, capturing steady-state phenomena still provides a good understanding of system properties (*4*).

As such, approaches often leave models with missing reactions, incorrect gene-reaction rules (*7*) or with pathways based on gap filling methods that add reactions that may not actually be catalysed by the organism in question (*33*). When attempting to understand specific natural phenomena, genome annotations are frequently updated and models are often corrected in an *ad hoc* manner.

**Figure 1:**
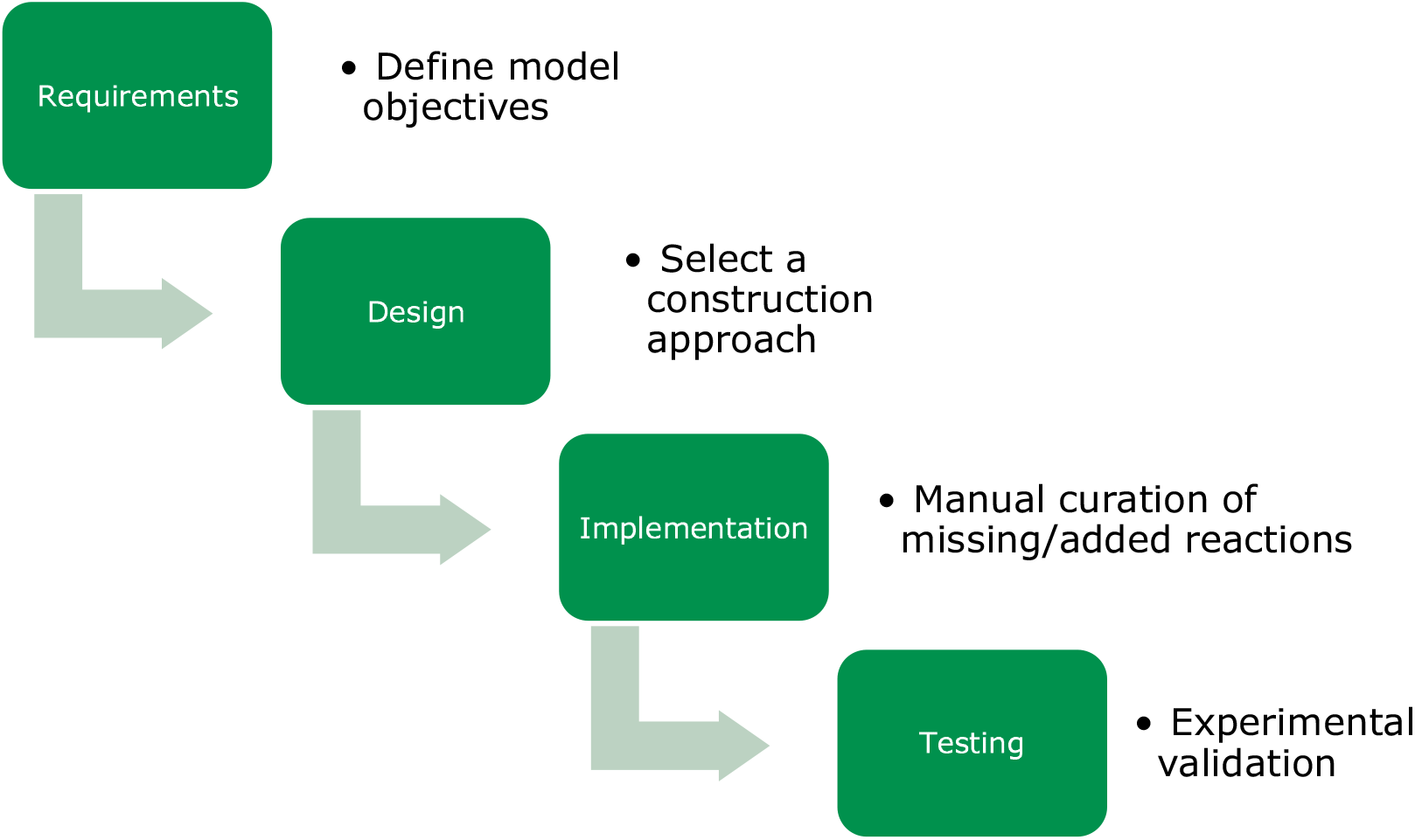
Conventional waterfall methodology for model development. Each process is considered to be an isolated aspect of model development.

Therefore, models undergo significant manual annotation and curation; a process which has a high chance of error. In this work, we advocate a test-driven approach to model development highlighted in Figure 2. Here, the model is changed to achieve research goals that are dynamic in response to the changes of a project. In order to meet this objective, validation criteria for a model such as growth conditions or the impact of gene knock outs, should be formally set. When a model is changed, all such validation criteria should be retested to ensure that models do not regress to previous states.

We feel that many of the current software tools for genome scale models do an excellent job of facilitating answers to crucial research and design questions. However, there is a major gap in terms of the reliability and re-usability of models due to a lack of standardisation and software tools to aid such processes. The following sections provide an overview of the *gsmodutils* software framework. *gsmodutils* aims to provide a basis for test-driven, version controlled agile model development. All software and packages are open source and is designed to be interoperable with platforms widely used in the domain of constraints based modelling.

**Figure 2:**
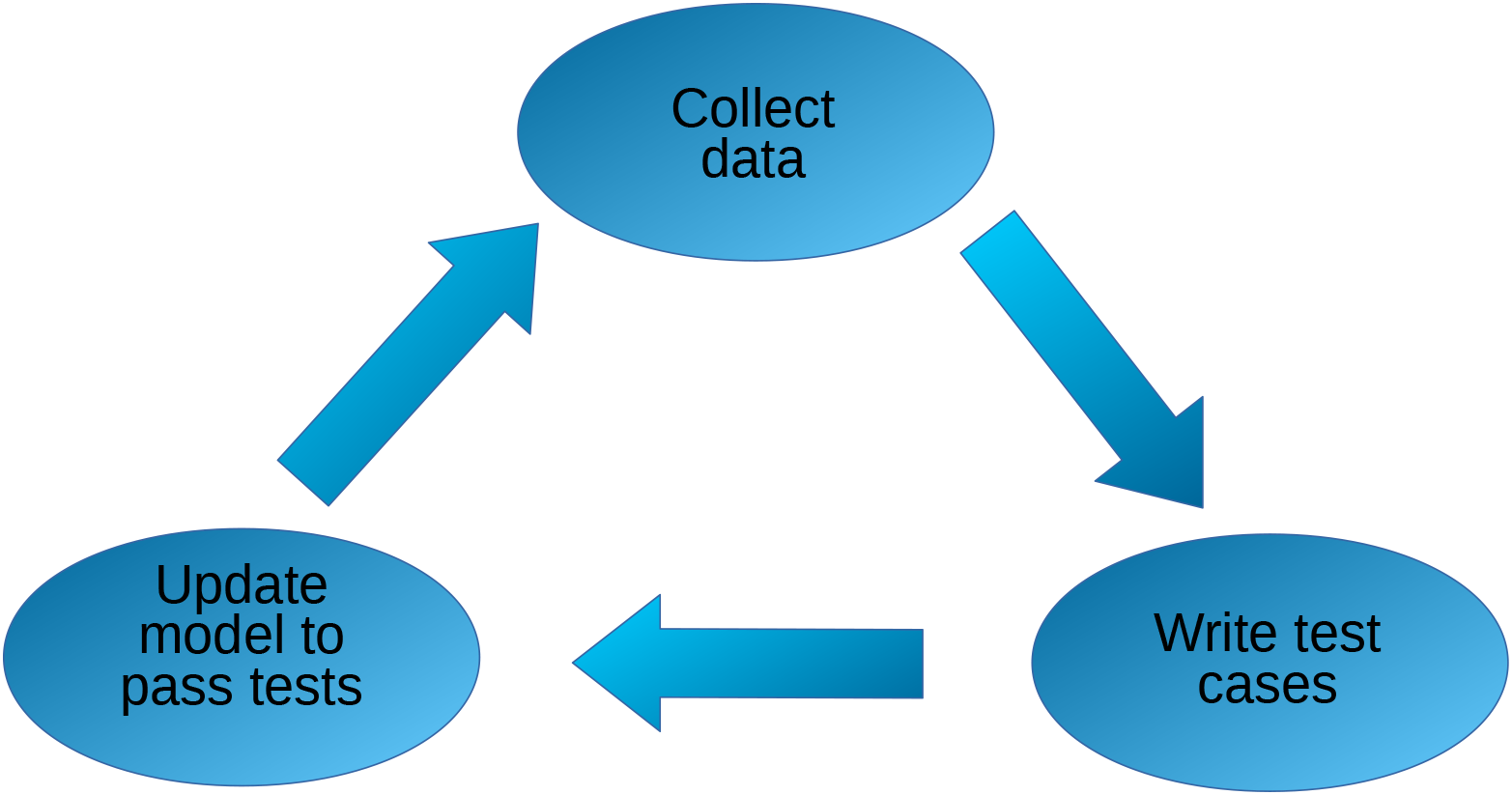
Workflow for model development proposed for *gsmodutils*. The objective of this approach is to simultaneously capture research questions, model validation criteria and minimise the impact of changes on previously completed models.

## 3 Software

### 3.1 Outline and features

Test-driven development is driven by the idea of clearly defined test cases written before significant changes are made to any underlying architecture. In the case of genome scale models, errors can easily occur as a product of human curation designed to better represent newly discovered aspects of metabolism.

By automatically integrating COBRApy (*13*) users can easily write convenient test cases following examples given in the user guide. A standard test case, ensuring that a given model grows on media is given in Figure 3. When a new model repository is created with the *gsmodutils* tool, a number of pre-written test cases are automatically added to a file. However, we stress that the vast majority of individual use cases for a model must be specific to a given biological problem.

The software provides a number of features such as import and export of models in different formats and the generation of test reports through use of the command line. The use of flat files enables easy integration with version control software such as git and mercurial. In addition, projects are easy to export using portable standardised docker images (*34*), the idea being to allow users to share models as quickly and easily as possible without concern for custom system configurations.

**Figure 3:**
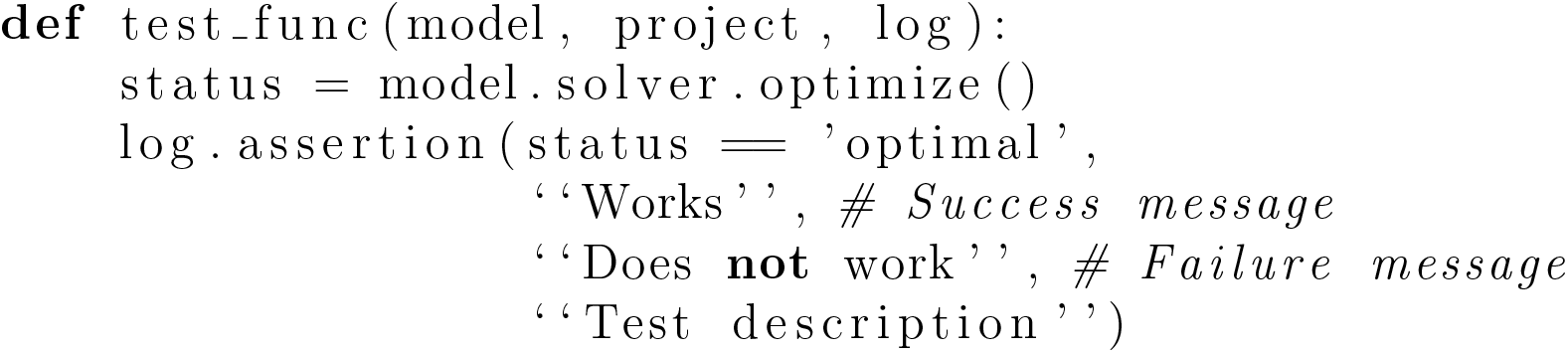
An example *gsmodutils* test case written in python

### 3.2 Strain designs

A core aspect behind the implementation of *gsmodutils* is the concept of a *design*, this encompasses a simple set of changes to a “wild-type” model that are required for analysis. For example a mutant strain that has undergone several gene deletions is not sufficiently different from the original wild-type to merit having its own genome scale model. However, it is often the case that such deletions are of scientific or industrial interest and, as such, the strain will be used in future work. Consequently, such designs are hereditary in nature. By taking the *model delta* between the constraints applied to an initial model and subsequent modifications, *gsmodutils* allows users to easily reuse and export models with this design. As designs inherit from a base model, future curation to a wild type model will automatically be included in the designs. Similarly, designs are self contained and will not interfere with one another allowing a form of project management. Figure 5 shows how this could work in a practical situation.

Designs of this nature can also be programmatic, allowing the implementation of features such as non-standard constraints that can be dynamically loaded. An example of this is shown in Figure 4. This example converts an existing model to one based on a Mixed Integer Linear Program (MILP), as the reaction names do not need to be specified, should reactions be added or removed the process will still apply. Alternative examples could include reductions of models through methods such as elementary flux modes or minimum cut sets, which can change dramatically with only small changes to stoichiometry. Furthermore, integrity of all strain designs is automatically tested as part of the default *gsmodutils* testing framework.

**Figure 4:**
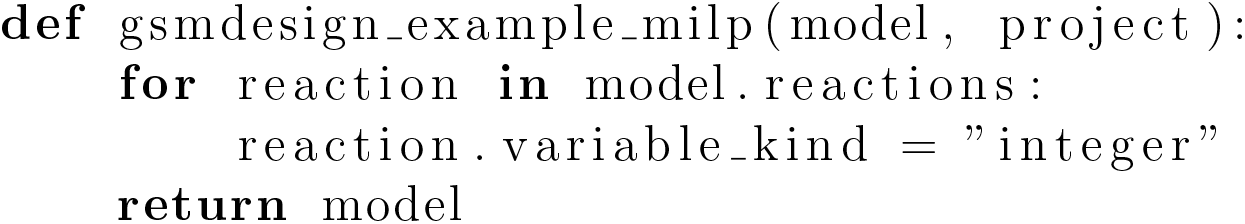
An example *gsmodutils* programmatic design written in python. This design converts reactions to integer type, allowing an MILP formation.

### 3.3 Development workflow

In this section we propose a method for the development of genome scale models that integrates *gsmodutils* with version control systems. Figure 3 highlights the notion of test cases, taken from test-driven development. The outlined workflow is that the user writes a formal test case for some modelling goal, perhaps driven by captured experimental data, that fits a specific form of validation criteria. We note that, in principle, this test case should be written before changes to a model.

**Figure 5:**
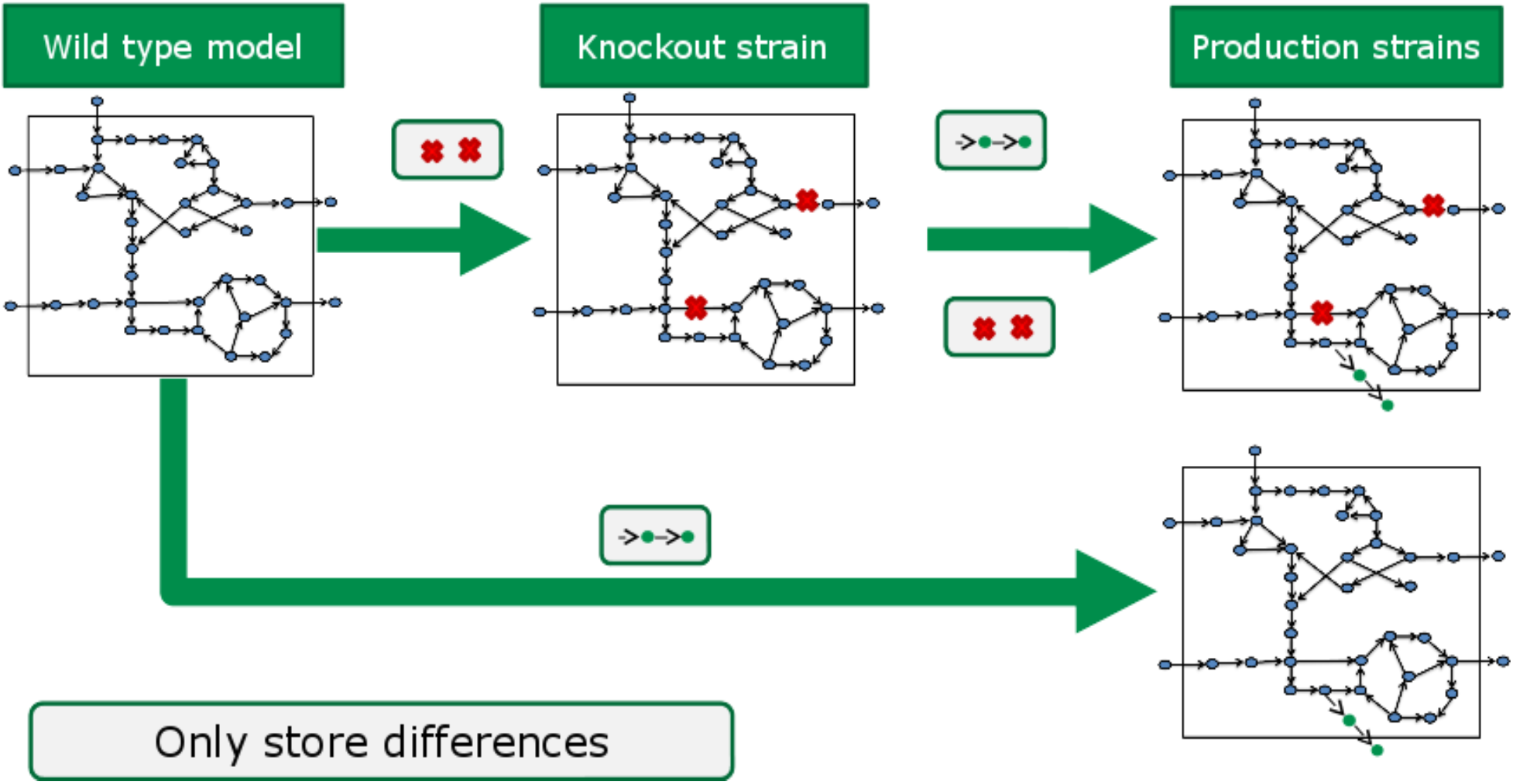
Examples of *gsmodutils* design inheritance. Each design stores the delta between the wild-type base model, any parents and the changes to constraints the design contains. In the example presented above a heterologous production pathway is combined with a reusable set of knock-outs. In practice, ideally, these designs should relate to real constructs and strains evaluated by experiment.

## 4 Case study usage Clostridium autoethanogenum

*Clostridium autoethanogenum* is a bacterial species used for the production of commodity chemicals at industrial scale (*35, 36, 37*). A new GSMM of *C. autoethanogenum*, ‘MetaCLAU’, has been analysed to improve this bioprocess ((*36*), submitted for peer review). In this section we describe how *gsmodutils* has been utilised to ensure that future versions of MetaCLAU will remain functionally relevant from the perspective of industrial biotechnology.

### 4.1 Scientific background and Model integration

*C. autoethanogenum* is a strictly anaerobic, acetogenic bacterium which naturally produces ethanol and trace amounts of *2,3-butanediol* (*2,3 BD*) from carbon monoxide and water (*35, 36, 38*). Since carbon monoxide is readily available in the form of industrial waste gas, and *2,3 BD* has a global market value of $43 billion (*39*), the optimisation of yields of *2,3 BD* from carbon monoxide is highly desirable in the context of industry (*36*). MetaCLAU was built using Pathway Tools (*40*) and ScrumPy (*9*), and is based on a manually annotated genome sequence of *C. autoethanogenum* (*41*). The resulting model consisted of 758 reactions, 773 metabolites and 518 genes. For full details of the model, see ((*36*). submitted for peer review). The model has been integrated with the *gsmodutils* modelling framework as a test-driven project. The following section details specific tests used to evaluate the model at each stage of its continued development.

### 4.2 Evaluation of model validation criteria

#### Energetic consistency

An important limitation of FBA is that optimal solutions may be thermodynamically infeasible if appropriate constraints are not applied (*42*). In order to identify these unwanted flux distributions and constrain the model such that they are not in the feasible solution space, a diagnostic FBA is applied with the following constraints: 1) All transport reactions are constrained to allow no uptake (lower bound = 0), and 2) the ATPase reaction is given a fixed flux of one. If a (non-zero) solution to this problem exists, it must contain a thermodynamic inconsistency, which can be dealt with by manual inspection of the solution and modification of one or more of the involved reactions (*42*).

#### Minimisation of flux based tests

One conventional approach in FBA is to set an optimisation criterion of minimising flux across enzymatic reactions with a fixed biomass constraint (*43*). The solution to this FBA problem represents minimal protein investment (*43*). Since execution of flux minimisation in COBRApy requires a model in which reversible reactions are split into two irreversible reactions, a *gsmodutils* design was created in which all reversible reactions are split in a programmatic manner. In the case of MetaCLAU, the flux minimal solution includes both ethanol and acetate production, which represents good qualitative agreement with experimental data ((*36*), under review). Since any changes to this predicted phenotype must be investigated, the flux minimisation analysis has been formulated as a *gsmodutils* test.

#### Product scans

Of interest to this project were changes in the product spectrum of *C. autoethanogenum* under conditions where the organism can and cannot produce molecular hydrogen (with carbon monoxide as sole carbon and energy source). The hypothesis tested in ((*36*), under review), was that in the case where hydrogen production infeasible, alternative electron sinks like lactate and *2,3 BD* would be produced. As in the previous case, the model-predicted behaviour showing both lactate and *2,3 BD* was deemed an important result, which model curators should be notified of if lost during model development. Thus the analysis was built into a *gsmodutils* test.

#### Lethal knockout mutants

The prediction of lethal single-gene KO mutants through FBA of a GSMM is useful in two ways: 1) the identification of essential genes is an important first step for metabolic engineering strategies, and 2) with the advent of high-throughput TraDIS gene-essentiality data sets (*44*), GSMMs can be validated by their ability to predict essential genes. Furthermore, any change in the set of essential genes (particularly an increase in the number of essentials) represents important information for metabolic engineering. For these reasons, a test has been built into the MetaCLAU project which enables the computation of the set of essential genes and their comparison with TraDIS data sets.

## 5 Related work

### Memote

(*45*) is tool set that has very similar ambitions to *gsmodutils*. It features a fully specified set of tests, including custom test cases and has strong version control integration with git. A core difference between these projects is that *gsmodutils* has a stronger focus on reducing the redundancy in model storage through the use of *design deltas*, as described above. Similarly, a core goal of *gsmodutils* is to allow easy import and export outside of the framework for compatibility with other modelling suites. It should be noted that, as memote is written in python, utilises COBRApy and, at the time of writing, it is fully compatible with *gsmodutils*.

### Cameo

(*14*) is a set of strain design utilities, such as genetic algorithms for finding optimal gene knock-out strategies, as well as other flux balance analysis utilities. Written in python and part of the COBRApy suite of applications, *gsmodutils* works well with cameo. Indeed, making a set of design changes can easily be integrated into *gsmodutils* through the python API.

### Model repositories

models are frequently shared, at the time of publication through services such as BiGG (*10*) and Biomodels (*46*). Whilst these repositories encourage the reuse of models and the reproducibility of *in silico* predictions they are not designed to improve collaboration. The software presented here is designed with the notion that genome scale models are never finished, *per se*, but under continuous development. The cornerstone of this is the use of test cases, which formalise modelling validation criteria.

## 6 Discussion

In order to facilitate the sharing and dissemination of high quality computational research, good standards and software are required (*47*). Naturally a great deal of effort has gone in to producing high quality systems and synthetic biology standards (*48, 49*). Furthermore, when research projects end it all to is common for important large models to be published and become relics lost within the literature, forgotten to all but the most dedicated of individuals. As GSMMs grow in terms of the metabolism the contain as well as the biological problems they are used to solve, problems with annotation and curation naturally accumulate as a product of human error. Software that facilitates actively improving how researchers develop and apply models to new phenomena is required.

We have presented a framework with a number of features taken from the software development world specifically designed to improve collaboration and minimise such error. However, it is important to stress the difference between defined behaviour expected from pre-written test cases and novel predictions made by a model. Indeed, a core objective of this framework is to ensure that good practices are followed in model development that help scientists to better trust the results discovered by their models. In an ideal world, we would envision a methodology such as ours becoming a pre-requisite for GSMMs to pass peer review.

As with most software development projects, *gsmodutils* will see expanded features. Initially this will include tighter integration with version control systems such as git and mercurial. Furthermore, the objective of the project is to cultivate collaboration by simplifying the process of distributing large models to different users. As a consequence, methods for sharing reusable docker containers will be investigated.

## Acknowledgement

We would like to thank the Oxford Brookes Cell Systems Modelling group for helpful discussions regarding this work and for scientific advice. This work was supported by the UK Biotechnology and Biological Sciences Research Council (BBSRC) and Engineering and Physical Sciences Research Council (EPSRC) grants BB/L013940/1, BB/K00283X/1 and BB/L502030/1.

